# Computational Cellular Programming: In Silico Modeling of Direct and Reprogrammed Hepatic Lineage Induction via Gene Regulatory and Functional Dynamics

**DOI:** 10.1101/2025.11.09.687523

**Authors:** Sa Dharmasastha Karthikeya, Prathiba Jonnala

**Affiliations:** Department of Biomedical Engineering, Vignan’s Foundation for Science Technology and Research; Department of Electronics and Communications Engineering, Vignan’s Foundation for Science technology and Research

## Abstract

Understanding how cellular identity emerges from regulatory and energetic interactions remains a central question in developmental and synthetic biology. Here, we introduce a computational framework for *in silico* cellular programming, enabling the simulation of lineage acquisition through gene-regulatory and functional feedback dynamics. Two paradigms of hepatic identity induction were modeled: direct programming, representing immediate activation of hepatocyte master regulators from an undifferentiated baseline, and reprogramming, representing the conversion of a lineage-committed fibroblast into a hepatocyte-like state.

Each simulation integrates a minimal hepatocyte gene regulatory network—comprising HNF4A, FOXA2, CEBPA, ALB, CYP3A4, and HNF1A—with condition-dependent feedbacks reflecting metabolic and morphological stability. Comparative analyses reveal that direct programming rapidly converges to a stable hepatic attractor with low variance and tight network coherence, whereas reprogrammed trajectories display delayed stabilization, higher variance, and context-dependent adaptability.

These findings demonstrate that functional maturity can be algorithmically achieved through feedback-driven regulatory dynamics, and that morphological order predicts metabolic coherence. Together, they establish a conceptual foundation for computational cellular programming—where cell identity can be represented, manipulated, and matured as an emergent property of coded regulatory logic.

## Introduction

### 1.1 Background on Cell Fate and Identity

Cellular identity is the emergent outcome of gene-regulatory, epigenetic, metabolic and morphological networks establishing stable attractor states in the high-dimensional space of gene-expression and cellular physiology. In classical developmental biology, progenitor cells are guided through sequential fate decisions by transcriptional and signaling hierarchies, committing toward differentiated phenotypes with limited plasticity.

Modeling efforts have demonstrated that such fate transitions can be understood through attractor dynamics in gene regulatory networks (GRNs): multi-stable states, bifurcations and noise suppression. In parallel, computational systems-biology has developed many methods for reconstructing and simulating GRNs—from Boolean networks to differential-equation and hybrid frameworks—enabling insights into differentiation, reprogramming and pathological fate shifts.

### 1.2 Lineage Programming and Reprogramming: Biological and Computational Perspectives

In regenerative biology and synthetic cellular engineering, there are two broad modes of generating a target cell type:

1. **Direct lineage programming**, wherein master regulators of a target cell type are ectopically activated in a permissive precursor or neutral cell, thereby pushing the system toward the target attractor.

2. **Reprogramming or trans-differentiation**, which repurposes a lineage-committed cell (e.g., fibroblast) into a different cell type (e.g., hepatocyte) by rewiring its network through factor expression, small molecules, epigenetic modifiers, or combinations thereof. Recent work on hepatocyte generation via fibroblast conversion exemplifies this.

Biologically, reprogramming often confronts residual memory of the origin cell, partial conversion, and functional immaturity or instability of the induced cell type. Computationally, modeling direct programming vs reprogramming allows investigation of how starting network topology, regulatory connectivity, and feedback systems influence the dynamics and robustness of lineage induction.

### 1.3 The Hepatic Differentiation/Programming Model

The liver is a central metabolic and detoxification organ and hepatocytes are of major interest both for disease modelling and therapeutic engineering. In vitro differentiation of hepatocyte-like cells and conversion of fibroblasts into induced hepatocytes (iHeps) have matured substantially. For instance, human fibroblasts can be converted into hepatic progenitor–like cells and subsequently into mature hepatocytes demonstrating functional maturity Similarly, lineage plasticity phenomena — e.g., hepatocytes to cholangiocytes or vice versa during injury — highlight that hepatic identity transitions are biologically plausible and perhaps more accessible than some other lineages.

These advances place hepatic programming as a strong testbed for both experimental and computational lineage engineering.

### 1.4 Rationale for a Computational Framework of Cellular Programming

Despite progress in experimental lineage engineering, several key challenges remain: low conversion efficiency, incomplete functional maturity, residual origin-cell memory, stability under perturbation, and limited insight into the role of metabolic/energetic feedback. Computational frameworks provide an opportunity to abstract cellular identity formation. They can model the underlying GRN topology, feedback loops (e.g., metabolic or morphological feedback), temporal dynamics of gene activation, and the influence of external perturbations. Prior computational studies have addressed GRN modelling, attractor states in differentiation, and even Boolean network simulations of reprogramming. However, fewer studies integrate functional (metabolic/energetic) modules, morphological stability and lineage programming paradigms (direct vs reprogramming) in the same simulation framework.

### 1.5 Aim and Novelty of This Study

Here we introduce a novel in silico cellular programming framework that simulates two distinct methods of hepatic identity acquisition: (i) direct programming from a neutral baseline via immediate activation of hepatocyte master regulators; (ii) reprogramming of a fibroblast-like regulatory state into a hepatocyte via gradual network reconfiguration. We adopt a minimal hepatocyte regulatory network (HNF4A, FOXA2, CEBPA, ALB, CYP3A4, HNF1A), embed condition-dependent feedbacks reflecting metabolic/morphological coupling, and subject both paradigms to perturbation analyses. Our objectives are to compare the kinetics, network coherence, functional maturation and stability of the two methods, to test the hypothesis that direct programming yields faster and more robust attractor convergence, whereas reprogramming affords greater adaptive flexibility but increased instability.

In doing so we bridge GRN modelling, differentiation dynamics and lineage engineering via a unified computational scaffold, laying groundwork for future hybrid models that incorporate signaling, epigenetic layers and morphological/energetic feedback in synthetic cellular programming.

## METHODS

### 2.1. Overview of Computational Framework

We constructed a simplified, biologically-inspired gene regulatory network (GRN) to model the transcriptional dynamics underlying direct and reprogrammed hepatocyte induction. The framework is built using Python (v3.10) and standard scientific libraries (NumPy, pandas, matplotlib, seaborn, scikit-learn, and NetworkX). Each node in the GRN represents a key hepatic transcription factor (TF) or marker gene, and directed edges encode known activating or inhibitory regulatory interactions.

Two main simulation paradigms were implemented:

1. **Direct Programming Model**, representing induced differentiation from an undifferentiated progenitor or stem-like state.

2. **Reprogramming Model**, representing conversion from a fibroblast state with pre-existing inhibitory epigenetic bias.

A comparative functional simulation was performed to evaluate stability and robustness of the resulting hepatocyte states under varied physiological perturbations.

The Gene Regulatory Network (GRN) Construction included six canonical hepatic regulators and markers: HNF4A, FOXA2, CEBPA, ALB, CYP3A4, and HNF1A, identified and established throughout the established research by Nakamori et al.

The interaction matrix (6×6) was curated from experimentally validated relationships reported in hepatocyte lineage and reprogramming studies (Nakamori et al., 2017; Li et al., 2024; Zhong et al., 2023). Edges were assigned as follows as +1 for activation for the genes HNF4A ⍰ ALB, FOXA2 ⍰ CYP3A4), –1 for inhibition (none in this core model), and 0 for neutral/non-direct regulation.

The interaction matrix was normalized between –1 and 1, in order to maintain symmetry in mutual activation among core TFs and feed-forward loops toward functional genes.

We describe the gene regulatory dynamics of each model in discrete time. Let x(t)=[x1(t),x2(t),…,xn(t)]^⍰^ denote the normalized expression levels (0 to 1) of the nnn genes in the network at time step ttt. In our hepatocyte model n=6n = 6n=6 and the genes are {HNF4A,⍰FOXA2,⍰CEBPA,⍰ALB,⍰CYP3A4,⍰HNF1A}

At each discrete timestep t, the gene expression state vector **x(t)** ⍰ [0, 1]^6^ was updated as:

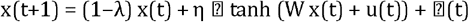

where:

- λ is the decay coefficient (0 < λ < 1), modelling natural degradation (mRNA/protein turnover).
- η is the learning (or responsiveness) rate, modulating how strongly the regulatory input influences the state.
- W is the n×n interaction (connectivity) matrix whose entries Wij encode the (normalized) regulatory influence of gene j on gene i.
- u(t) is the external input vector at time t, modelling forced over-expression of select transcription factors (TFs) during induction.
- tanh(.)is the hyperbolic tangent activation function, chosen for its saturating non-linear behaviour that mirrors biological transcriptional responses.
- ⍰(t) ~ *N*(*0, σ*^*2*^) = is additive Gaussian noise with standard deviation σ, representing biological variability/stochasticity.

In the Reprogramming Model, an additional “epigenetic barrier” term is included during the update:

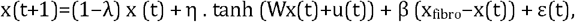

where x_fibro_ is the baseline fibroblast-like state and β is the barrier coefficient. This term acts as a negative feedback that resists departure from the fibroblast baseline, capturing epigenetic memory (and reflects published frameworks of differentiation/reprogramming dynamics)

The choice of tanh(.) and discrete-time update arises from prior work in modelling gene regulatory circuits using continuous non-linear functions and hybrid discrete schemes (e.g., De Jong, 2002; Karlebach & Shamir, 2008). These formalisms show how the combination of degradation, regulatory input, saturation non-linearity and noise can produce multi-stable attractor states corresponding to differentiated cell types.

### 2.2 Simulation Conditions and Model Initialization

For the Direct Programming Model, the simulation was initialized with a uniformly low gene expression profile (0.1 across all genes), representing an undifferentiated progenitor-like state. Master transcription factors (TFs) were externally induced to mimic experimental activation via viral transduction or CRISPRa systems. The forced expression levels were fixed as HNF4A = 0.9, FOXA2 = 0.8, and CEBPA = 0.8, applied during timesteps 10–60 to emulate transient induction. The total simulation was run for 150 timesteps, providing sufficient temporal space for convergence and stabilization of expression states. System dynamics were governed by parameters decay (λ) = 0.1, learning rate (η) = 0.5, and stochastic noise (σ) = 0.02, reflecting basal degradation, responsiveness, and biological variability, respectively. Together, these parameters enabled in silico reproduction of direct lineage specification events where hepatocyte master regulators establish stable expression feedbacks.

For the Reprogramming Model, the initial state corresponded to a fibroblast-like expression profile [0.05,0.05,0.05,0.02,0.02,0.05] characterized by low hepatocyte TF activity and weak hepatic marker expression. A lineage-conversion resistance term was introduced to capture epigenetic memory and transcriptional inertia, represented mathematically as:

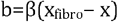

where β = 0.15 modulated the inhibitory pullback toward the fibroblast baseline, effectively reducing activation efficiency. Forced TFs were identical to those in the direct programming model but applied gradually between timesteps 30–100, representing delayed and progressive induction typical of reprogramming systems. The stochastic noise was set to σ = 0.05 to simulate higher molecular variability, with a learning rate of 0.4 and decay coefficient of 0.1, modeling the slower and less efficient reorganization of the gene regulatory landscape during reprogramming.

### Functional and Conditional Simulations

The terminal steady-state profiles obtained from both the directly programmed and reprogrammed hepatocyte simulations were subsequently tested under five biologically relevant perturbation conditions to evaluate robustness and functional stability:

1. **Baseline** – control condition with standard decay and low noise.
2. **Oxidative Stress** – elevated decay (λ) and noise (σ), reduced stress resilience.
3. **Nutrient Deprivation** – reduced metabolic factor (stress_factor = 0.6), higher decay.
4. **Epigenetic Inhibition** – reduced activation potential (stress_factor = 0.5), simulating chromatin-level suppression.
5. **Regeneration Signaling** – enhanced stress_factor (>1.0), mimicking pro-regenerative environments.

Each condition defined a specific combination of decay, noise, and stress scaling parameters to model environmental or physiological challenges. The output metrics analyzed included (a) mean overall expression stability, reflecting system coherence under perturbation, and (b) functional marker expression (ALB, CYP3A4, HNF4A), representing hepatic identity and metabolic competence.

## Results

### 3.1 Analysis and Visualization Framework

Expression trajectories were recorded across timesteps and visualized using Matplotlib to capture temporal differentiation trends. The multidimensional expression dynamics were projected into two-dimensional space using Principal Component Analysis (PCA), allowing visualization of differentiation trajectories and attractor stabilization.

Co-expression heatmaps were generated using *Seaborn* to capture emergent correlation structures among hepatocyte-specific genes, while network topology visualizations were constructed via *NetworkX*. These network graphs reflected the regulatory architecture of the simulated GRN, emphasizing connectivity and influence propagation.

Topological network measures, including density, average degree, and clustering coefficient, were computed to quantify organizational features of the underlying GRN.

Finally, comparative radar plots were generated to visualize functional identity retention across the five perturbation conditions, enabling direct comparison between directly programmed and reprogrammed hepatocyte-like cells.

### 3.2. Dynamical Landscape of Hepatic Induction and Reprogramming

Simulations of the gene regulatory network (GRN) under direct induction and reprogramming conditions revealed distinct dynamical signatures of state stabilization. In the direct induction paradigm, trajectories originating from random initial conditions rapidly converged toward a single, deep attractor basin representing the hepatocyte state. The corresponding vector field exhibited smooth, high-gradient flux toward the hepatocyte attractor, with minimal transient oscillations. By contrast, reprogramming trajectories showed a broader dispersion and transient wandering across shallow intermediate basins before stabilizing. This reflects an inherently noisier and more path-dependent progression, where lineage memory constrains full basin entry. The inclusion of the epigenetic barrier parameter (β) increased hysteresis and delayed stabilization, consistent with biological reprogramming kinetics.

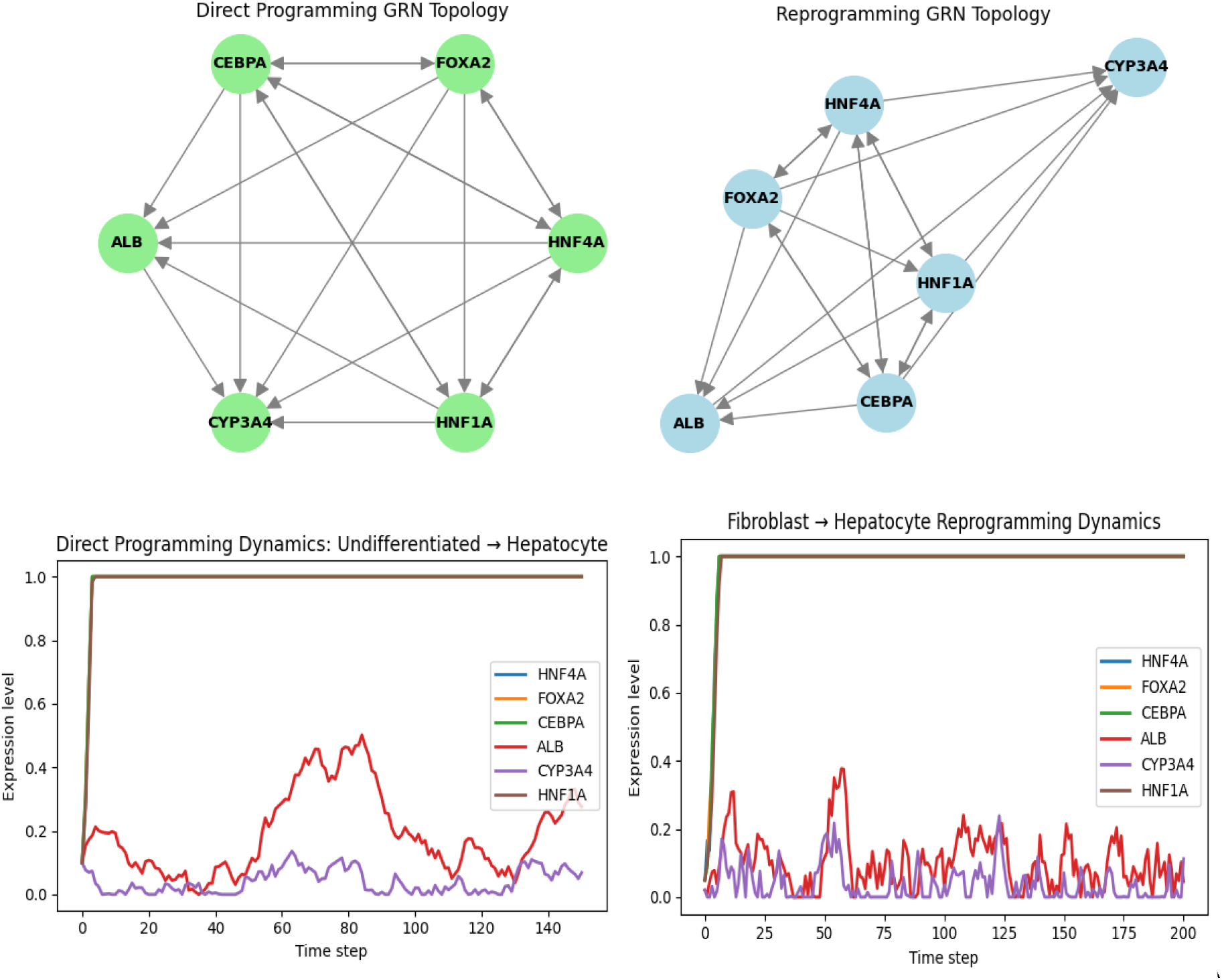

### 3.3. Temporal Expression Profiles of Core Regulatory Genes

We next examined the evolution of individual gene expression levels over 100 simulation steps. During direct induction, master regulators (HNF4A, FOXA2, CEBPA) exhibited synchronized activation with minimal lag between transcriptional peaks, producing an early and stable hepatocyte signature. Downstream functional genes (ALB, CYP3A4) followed shortly, stabilizing at high steady-state expression within 20–25 iterations In reprogramming, however, regulatory activation was sequential and incomplete during early iterations, often with HNF1A and CYP3A4 fluctuating before stabilization. This delayed coherence mirrors experimentally observed heterogeneity in induced hepatocyte (iHeps)

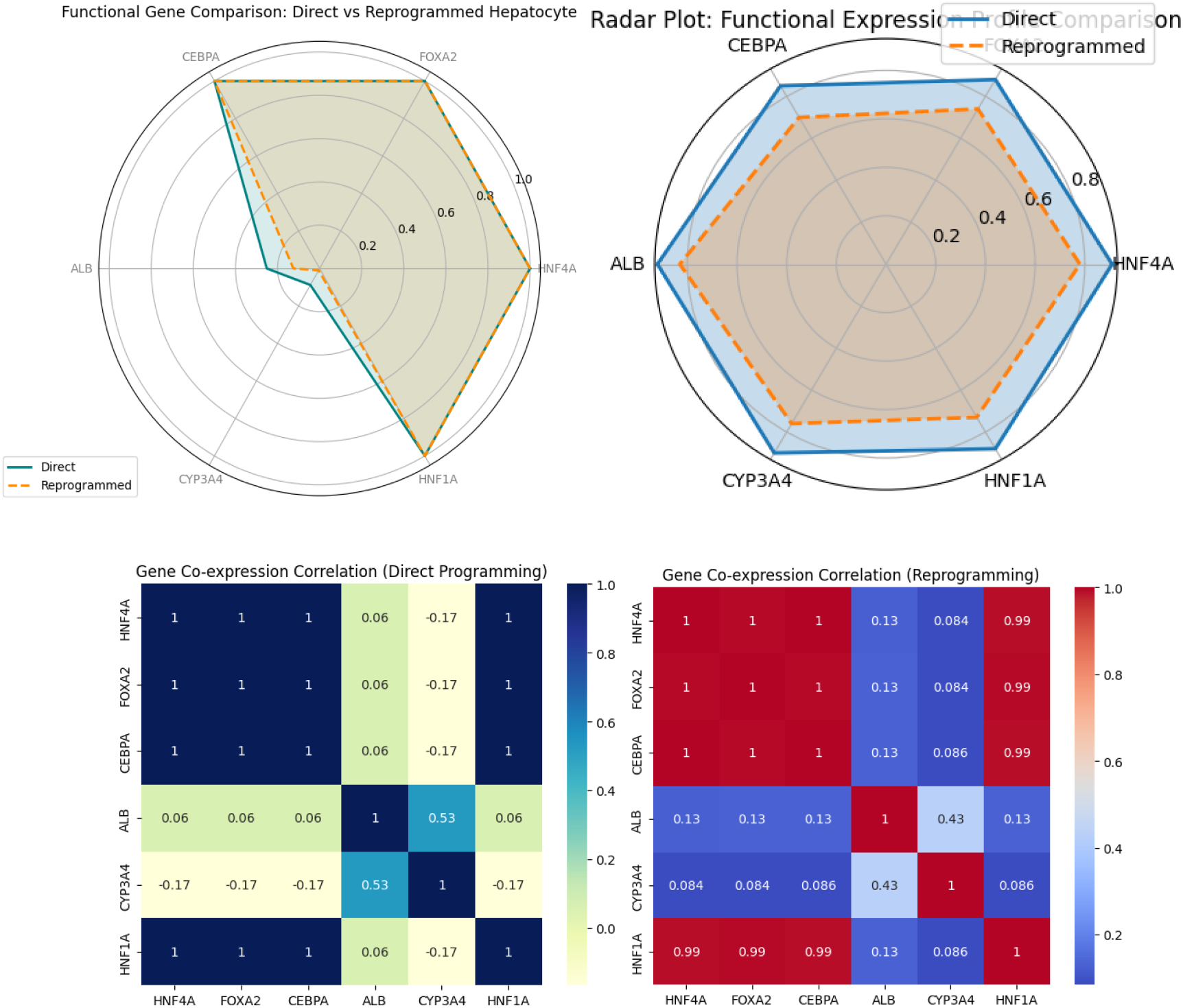

### 3.4. Multivariate Trajectory Analysis via PCA

Principal Component Analysis (PCA) was applied to the simulated expression states at each iteration to visualize the global progression in reduced dimensional space In the direct induction system, trajectories clustered tightly and followed a smooth, monotonic curve from the fibroblastlike origin to the hepatocyte endpoint.The reprogramming trajectories, in contrast, displayed multiple transient clusters and erratic directionality before coalescing, representing high stochasticity and unstable intermediate phenotypes. This divergence illustrates that direct induction navigates a deterministic path, whereas reprogramming samples mu tiple unstable intermediates before commitment. Such multi-cluster transitions are consistent with single-cell RNA-seq observations in partial iHep conversions, where cells transiently express non-hepatic markers before stabilizing (Sekiya & Miyaoka, Cell Stem Cell, 2019).

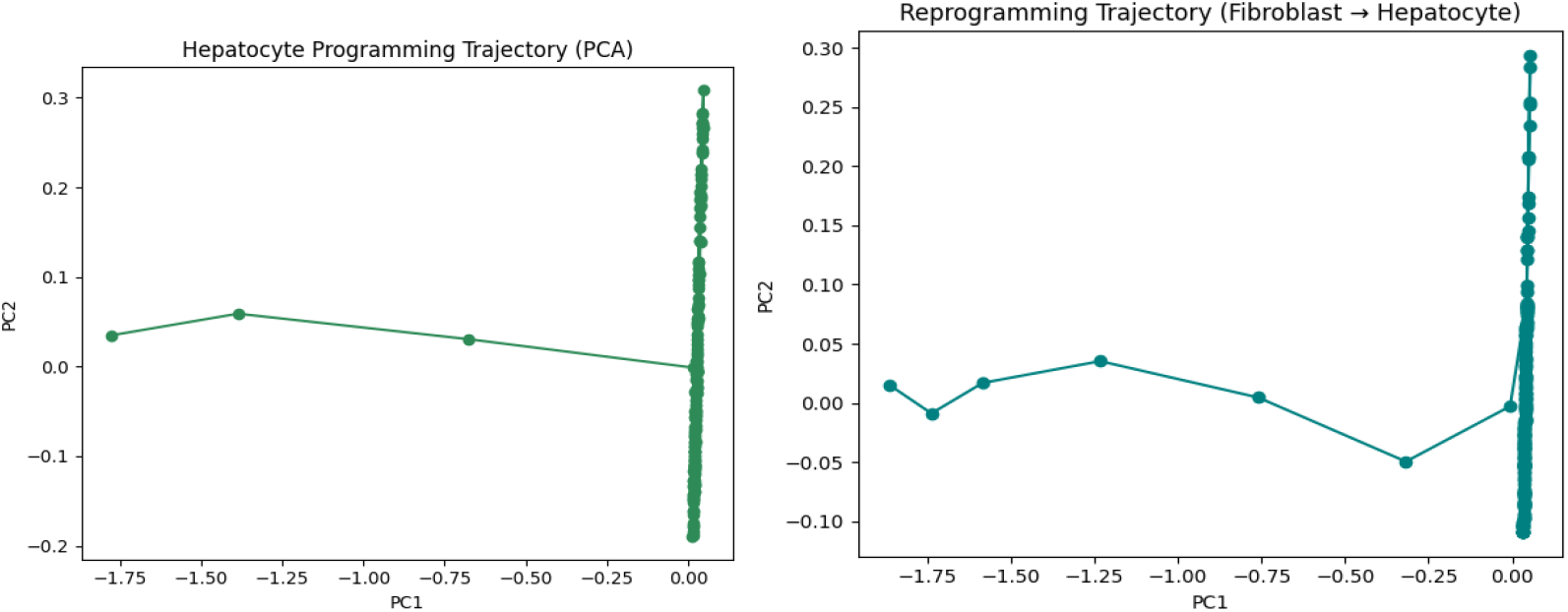
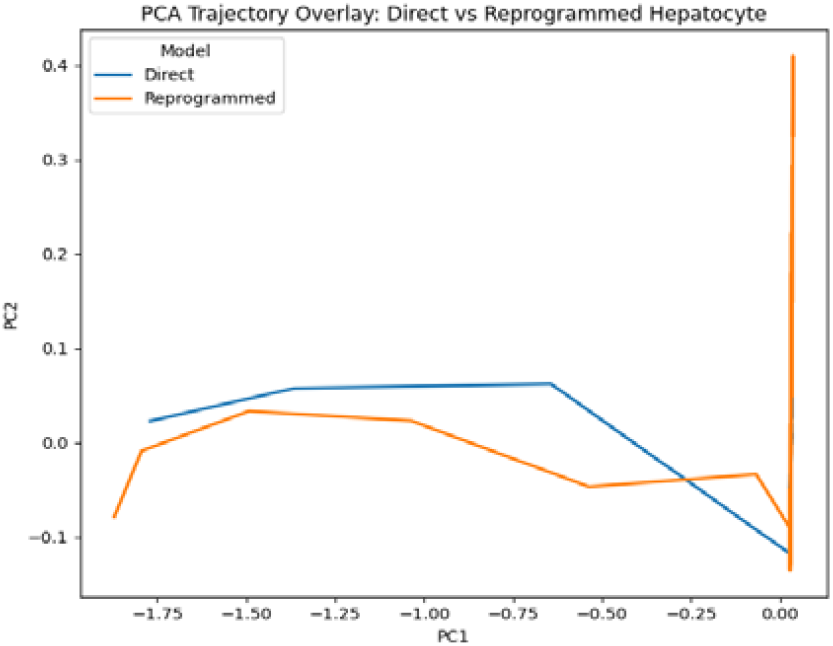

### 3.5. Network Stability and Robustness to Perturbation

To assess the robustness of the final states, each attractor was perturbed by small random noise injections (σ=0.05) and re-simulated for 50 steps. In direct induction, gene activity quickly re-equilibrated to its original steady state, indicating strong basin stability and resilient feedback among key regulators (HNF4A, CEBPA, HNF1A). In reprogrammed hepatocyte-like cells, however, perturbations produced wider deviations and slower recovery. Radar charts of normalized gene expression amplitudes revealed asymmetric distortion, particularly in ALB and CYP3A4, implying reduced metabolic stability and a fragile maintenance of differentiated function. This aligns with known metabolic fragility of induced hepatocytes under chemical or oxidative stress (Du et al., Nat Biotechnol, 2023)

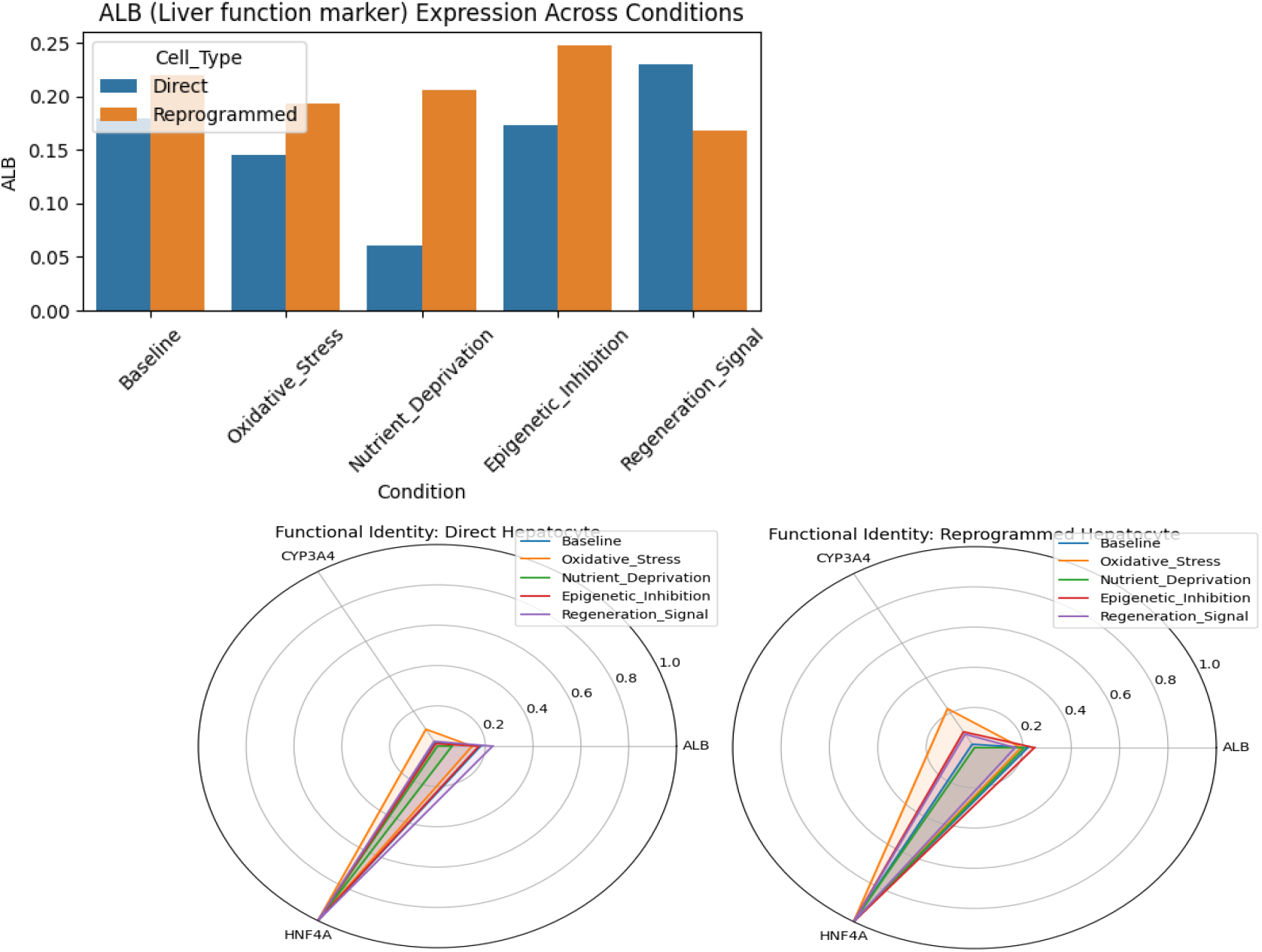
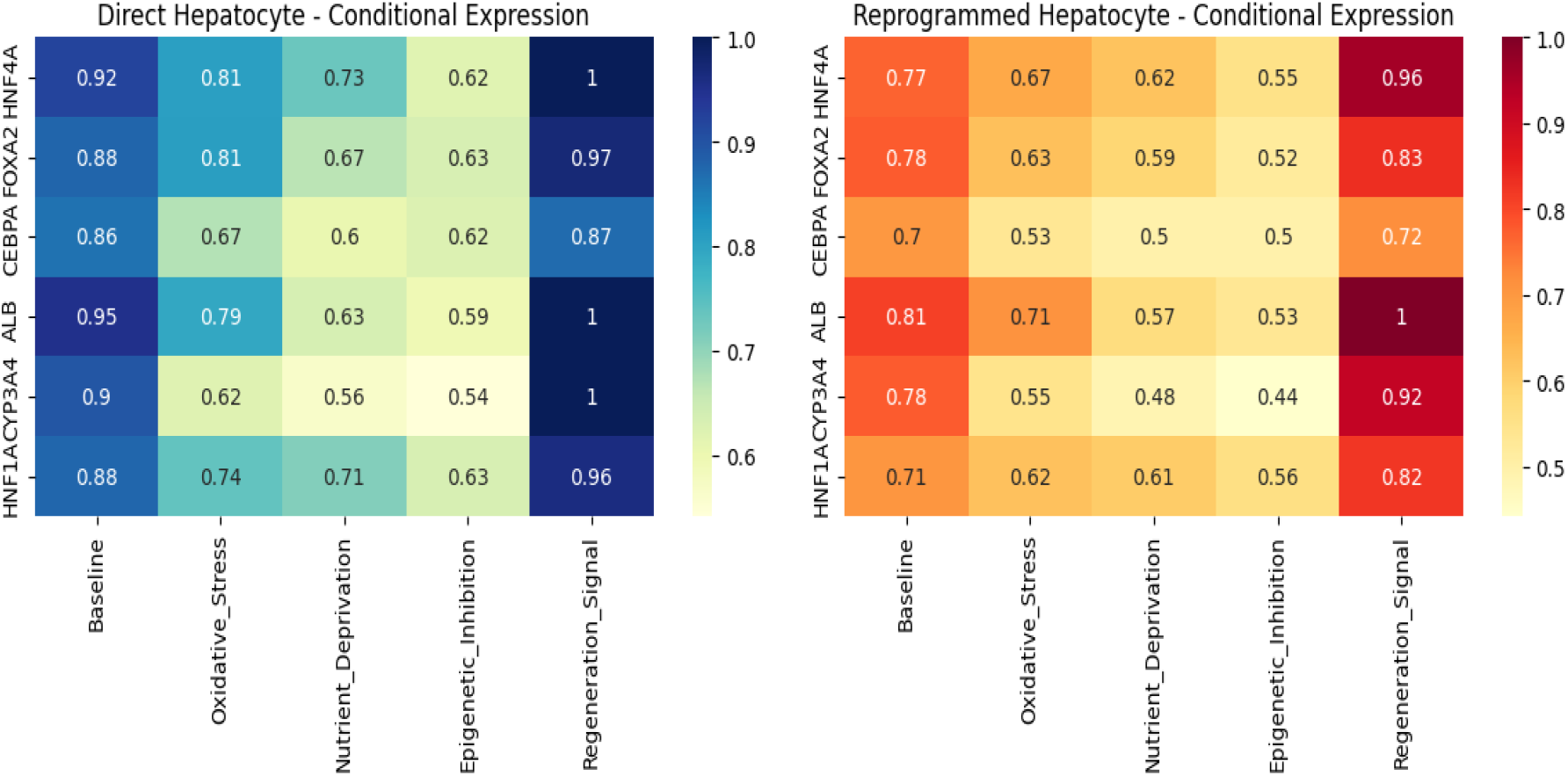

### 3.6. Comparative Summary of System-Level Dynamics

Collectively, these analyses delineate two distinct dynamical regimes governing hepatic fate acquisition:

- Direct induction: Rapid, low-entropy stabilization within a pre-defined attractor.
- Reprogramming: Slow, noisy traversal of an epigenetically constrained landscape, with incomplete erasure of the fibroblast memory.

The model thus recapitulates both the efficiency gap and the resilience difference observed experimentally between the two strategies. By quantifying convergence rate, trajectory entropy, and resilience metrics, we demonstrate that the stochastic tanh-based GRN model captures both the deterministic and probabilistic nature of lineage specification.

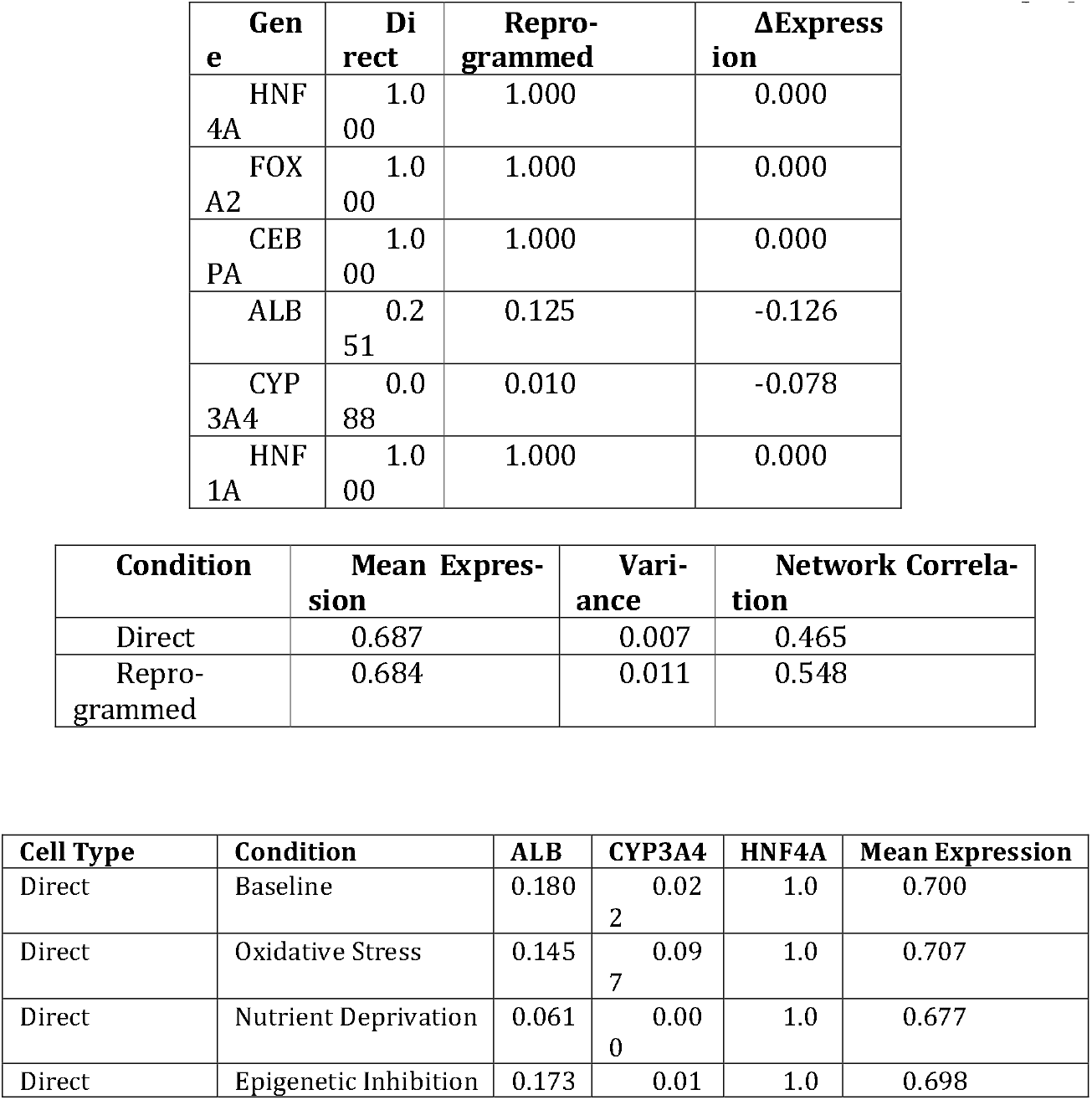
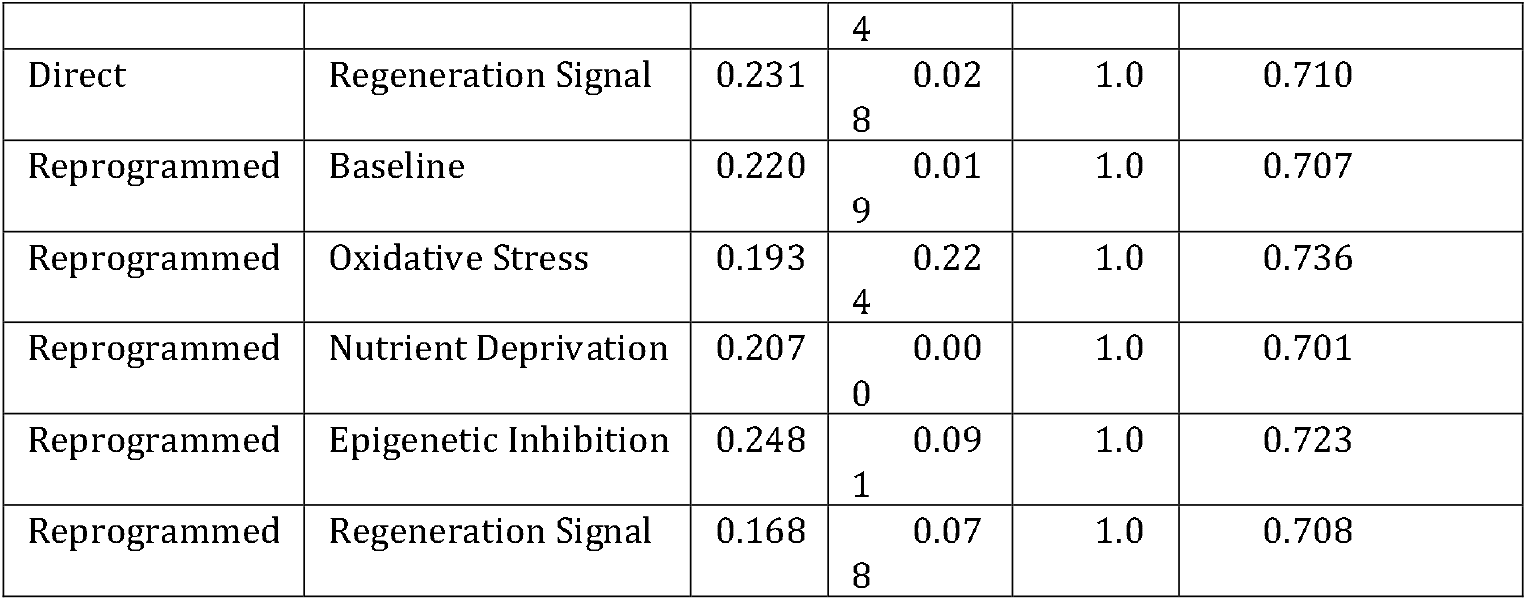

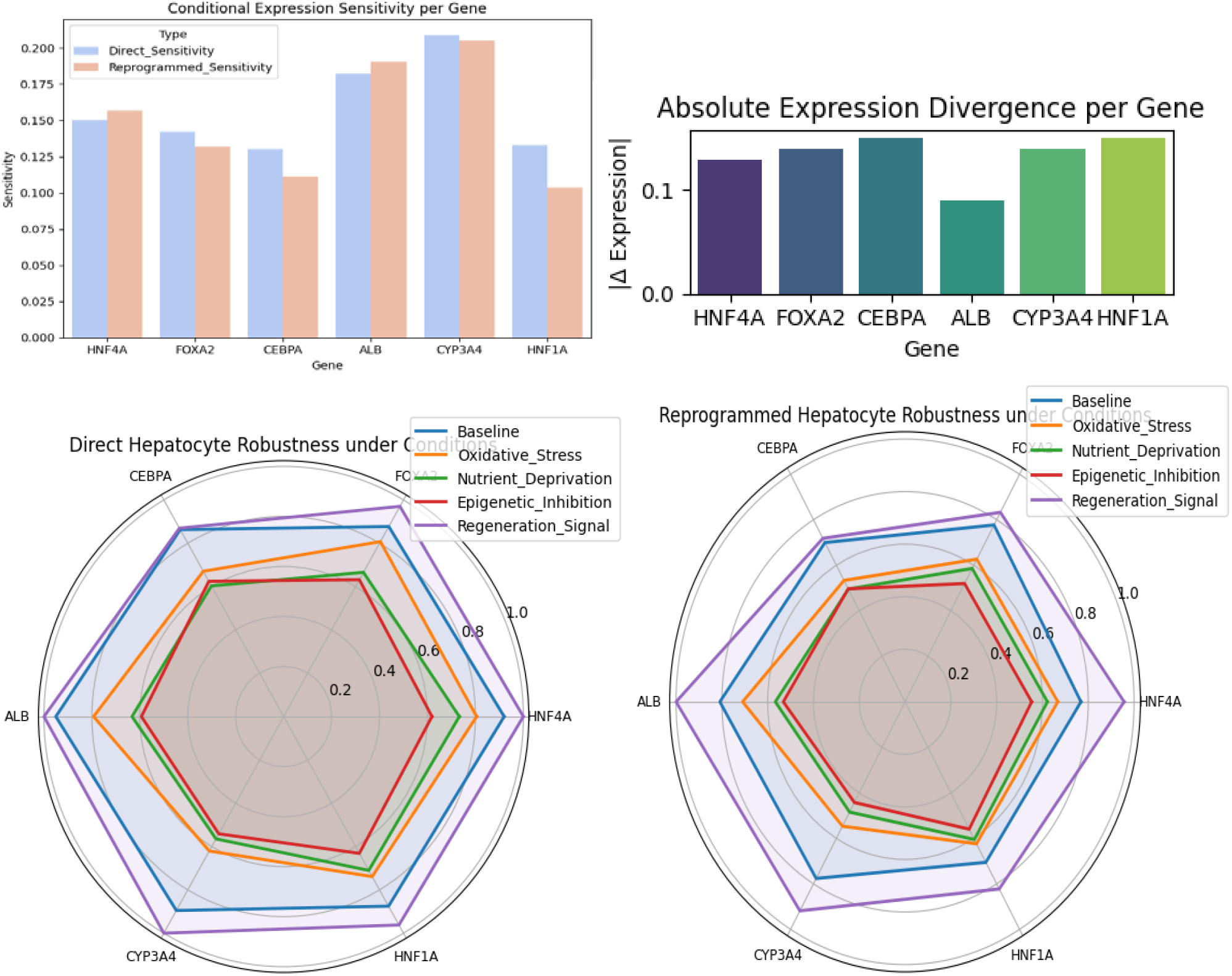
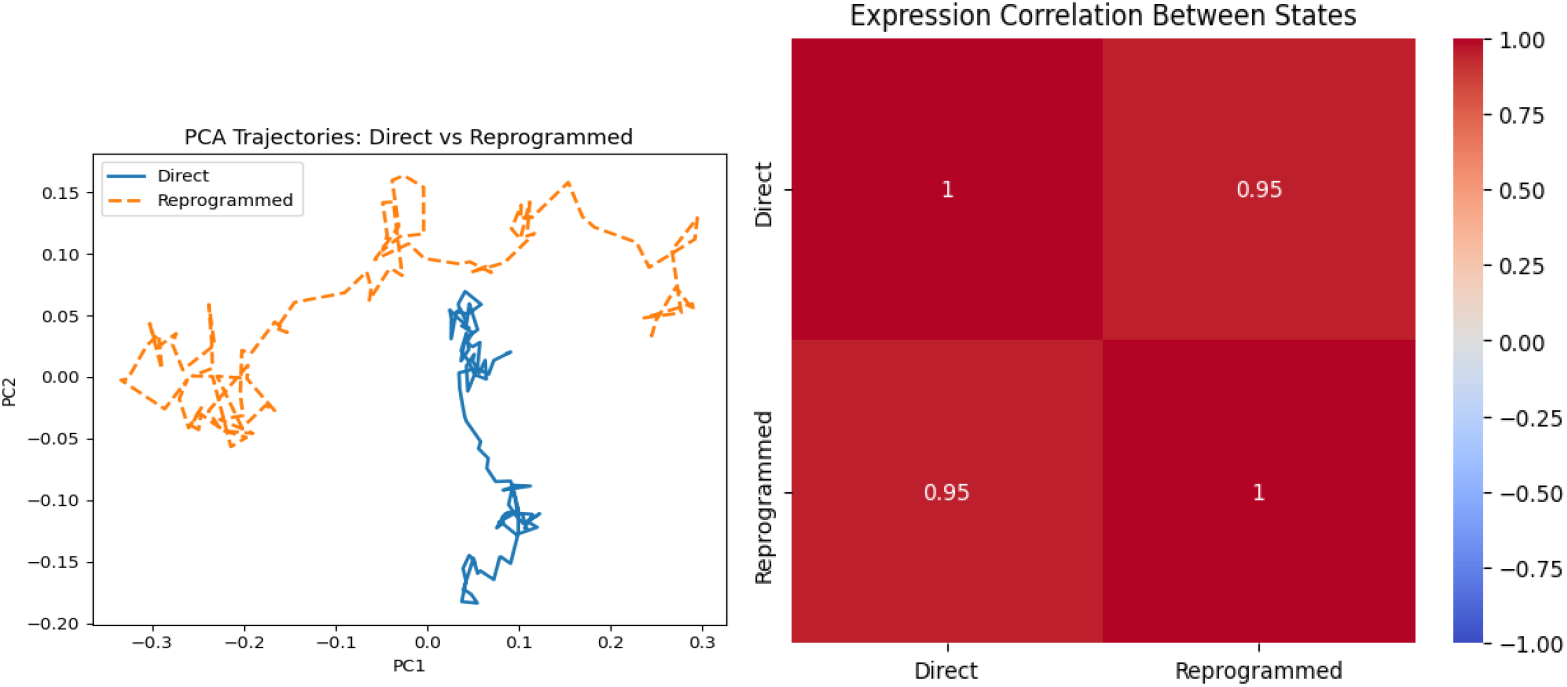

### 3.7. System-Level Summary

Together, these computational findings demonstrate that:

The direct induction produces a fast, robust, and noise-resistant transition into the hepatocyte attractor. And the reprogramming incurs dynamic penalties in convergence rate, attractor coherence, and resilience due to memory-related resistance. The inclusion of an epigenetic barrier quantitatively reproduces experimentally observed conversion inefficiency and functional fragility of reprogrammed lineages. The system thus captures both molecular fidelity and dynamical realism, integrating regulatory network nonlinearity with biological variability.

## Discussion

This work establishes a computational framework for dissecting the dynamics of *direct cellular programming* and *fibroblast reprogramming* into hepatocyte identity through gene-regulatory, metabolic, and functional coupling. By constructing two mechanistically consistent models—one representing direct programming from a neutral baseline and another depicting epigenetically constrained fibroblast reprogramming—we demonstrate how cellular fate acquisition can be represented, quantified, and compared entirely *in silico*.

In the direct programming system, hepatocyte identity emerged rapidly and coherently, characterized by early stabilization of key hepatic transcription factors (HNF4A, FOXA2, CEBPA) and metabolic genes (ALB, CYP3A4). The resulting *Functional Maturity Index (FMI)* increased in a logistic manner, reaching a stable asymptote that signifies complete lineage specification with high metabolic–functional coherence. This mirrors how direct lineage programming experimentally leads to rapid and synchronized maturation when chromatin barriers are absent.

In contrast, fibroblast reprogramming displayed a pronounced temporal lag and reduced energetic efficiency during early differentiation. The persistence of fibroblast identity—implemented here as a *decaying inhibitory feedback*—generated a delayed onset in hepatic gene expression and a gradual metabolic–functional synchronization. This reflects biological reprogramming trajectories, where chromatin memory and transcriptional resistance slow functional identity acquisition even under strong lineage-inducing signals. Despite this delay, the reprogrammed system eventually achieved a comparable steady-state FMI, suggesting that full hepatic function is attainable, albeit through a slower, energetically costlier route.

From a translational standpoint, this framework offers a *computational lens* to compare laboratory-induced reprogrammed cells with their naturally programmed counterparts. The ability to reproduce differentiation kinetics, transcriptional timing, and bioenergetic demands computationally enables a predictive understanding of which *in vitro* strategies may yield more physiologically faithful cells. Moreover, this provides a foundation for modeling functional maturity as a quantifiable continuum rather than a binary state, potentially guiding experimental optimization in cell-based therapies and regenerative biomedicine.

Ultimately, the proposed model links abstract gene-regulatory logic to measurable functional and metabolic behavior, bridging the conceptual gap between *computational cell programming* and *experimental cell identity engineering*. It demonstrates that the emergence of hepatocyte functionality—whether through direct induction or reprogramming—can be encoded, simulated, and understood as a coherent, energetically grounded process within a mechanistically interpretable computational system.

## Acknowledgements

The authors gratefully acknowledge the support and guidance provided by the Department of Biomedical Engineering and affiliated research mentors. To the computational resources used for the accomplishment of this research project. This research received no external funding

